# *“mir152* hypomethylation, potentially triggered by embryonic hypoxia, as a common mechanism for non-syndromic cleft lip/palate”

**DOI:** 10.1101/850016

**Authors:** Lucas Alvizi, Luciano Abreu Brito, Bárbara Bischain, Camila Bassi Fernandes da Silva, Sofia Ligia Guimaraes Ramos, Gerson Shigeru Kobayashi, Jaqueline Wang, Maria Rita Passos-Bueno

## Abstract

Non-syndromic cleft lip/palate (NSCLP), the most common human craniofacial malformations, is a complex disorder given its genetic heterogeneity and multifactorial component revealed by genetic, epidemiological and epigenetic findings. Association of epigenetic variations with NSCLP has been made, however still of little functional investigation. Here we combined a reanalysis of NSCLP methylome data with genetic analysis and used both *in vitro* and *in vivo* approaches to dissect the functional effects of epigenetic changes. We found a frequent differentially methylated region in *mir152*, hypomethylated in NSCLP cohorts (21-26%), leading to *mir152* overexpression. *In vivo* analysis using zebrafish embryos revealed that *mir152* upregulation leads to craniofacial impairment analogue to palatal defects. Also, we demonstrated that zebrafish embryonic hypoxia leads to *mir152* upregulation combined with *mir152* hypomethylation and also analogue palatal alterations. We therefore suggest *mir152* hypomethylation, potentially induced by hypoxia in early development, as a novel and frequent predisposing factor to NSCLP.

## Introduction

Non-syndromic cleft lip/palate (NSCLP) is the most common craniofacial congenital malformation in humans, affecting 1:700 live-births worldwide, and follows a multifactorial model of inheritance ^1–3^. Genetic contribution to NSCLP has long been supported by several independent studies, which has shown heritability estimates as high as 78-91% in Asian, European and Brazilian populations ^4–6^. Genomic analyses have successfully revealed several at-risk common genetic variants, in distinct populations. Nevertheless, they confer a small risk, and explain 10-30% of the disease’s heritability ^7,8^. In addition, an increasing number of rare pathogenic variants has also been identified in families segregating NSCLP, but the extent of their contribution in overall NSCLP cases is uncertain; importantly, no shared prevalent genetic basis has been observed for these variants ^9–14^, except for mutations in the Epithelial Cadherin-p120-Catenin Complex, which are responsible for 2-14% of familial NSCLP cases ^15^. Given the lack of a common mechanism underlying a large proportion of cases, projections for strategies of prevention and development of predictive diagnostic tests in at-risk couples have been currently hindered.

In parallel with genetic studies, epidemiological studies have suggested the influence of several environmental factors predisposing to NSCLP ^16–23^. In this sense, recent progress on uncovering the epigenetic contribution to NSCLP have been made ^24–27^. Epigenetic variations (or epivariations) are dynamic, functional and inheritable covalent changes in DNA and/or chromatin associated proteins which do not alter DNA sequence, yet they can affect gene expression and contribute to phenotypic variability and disease ^28–33^. Association of genomic epivariations to phenotypes, so called Epigenome-wide association studies (EWAS), have been expanding the knowledge on phenotypic variability and disease molecular mechanisms for the past years ^30,31,34–41^. More recently, individual-specific methylome analysis has shed light on epigenetic variation relevant to disease, demonstrating how this approach can uncover molecular alteration for complex traits ^42^. Here, we attempted to identify both group and individual-specific methylation changes using previously published methylome data on NSCLP. We identified individual methylation changes in known NSCLP candidate regions and also *mir152* hypomethylation in 26% of our discovery cohort. This result was replicated in an independent cohort and validated through functional *in vitro* and *in vivo* assays. Finally, we demonstrated how hypoxia, a known environmental risk factor for NSCLP, can modulate such changes.

## Results

### mir152 is a frequent differentially methylated region in the Brazilian NSCLP cohort

We conducted differential methylation analysis at the gene level using the whole Brazilian NSCLP 450K dataset (66 NSCLP vs 59 controls ^24^, and looked for the top 5 DMRs ranked by RnBeads, which combines adjusted p-value to methylation difference and methylation quotient. Those top DMRs were, in order of ranking: top 1, an intronic region of *CROCC* at 1p36.13; top 2, an intronic region of *FAM49B* at 8q24.21; top 3, an intronic region of *NLK* at 17q11.2; top 4, a non-coding region comprising *mir152* at 17q21.32; and top 5, an exonic region of *PRAC2* and comprising *mir3185* also at 17q21.32 (Figure 1a; Supplementary Table 2). Among those genes, *mir152* (adjusted p-value= 8.20E-06, beta-difference = −0.04) was the only with enriched expression during palatal embryogenesis in human and mouse, according to Sysface (Systems tool for craniofacial expression-based gene discovery) online tool. Moreover, *mir152* has already been identified as a DMR during normal murine palatal development ^53^ and suggested as a central regulator of downstream mRNAs encoding proteins known to play pivotal roles in orofacial development ^54^, however still with no clear evidence of association with NSCLP. Concurrently, we also conducted a differential methylation analysis at the gene level comparing individually each one of the 66 NSCLP samples versus all 59 controls (“450k cohort”), looking for individual epivariation. We found a total of 6620 gene DMRs (average = 100.3 DMRs per sample) in all NSCLP samples with >5% methylation difference and adjusted p-value<0.05 (Supplementary Table 3).

**Figure 1:**
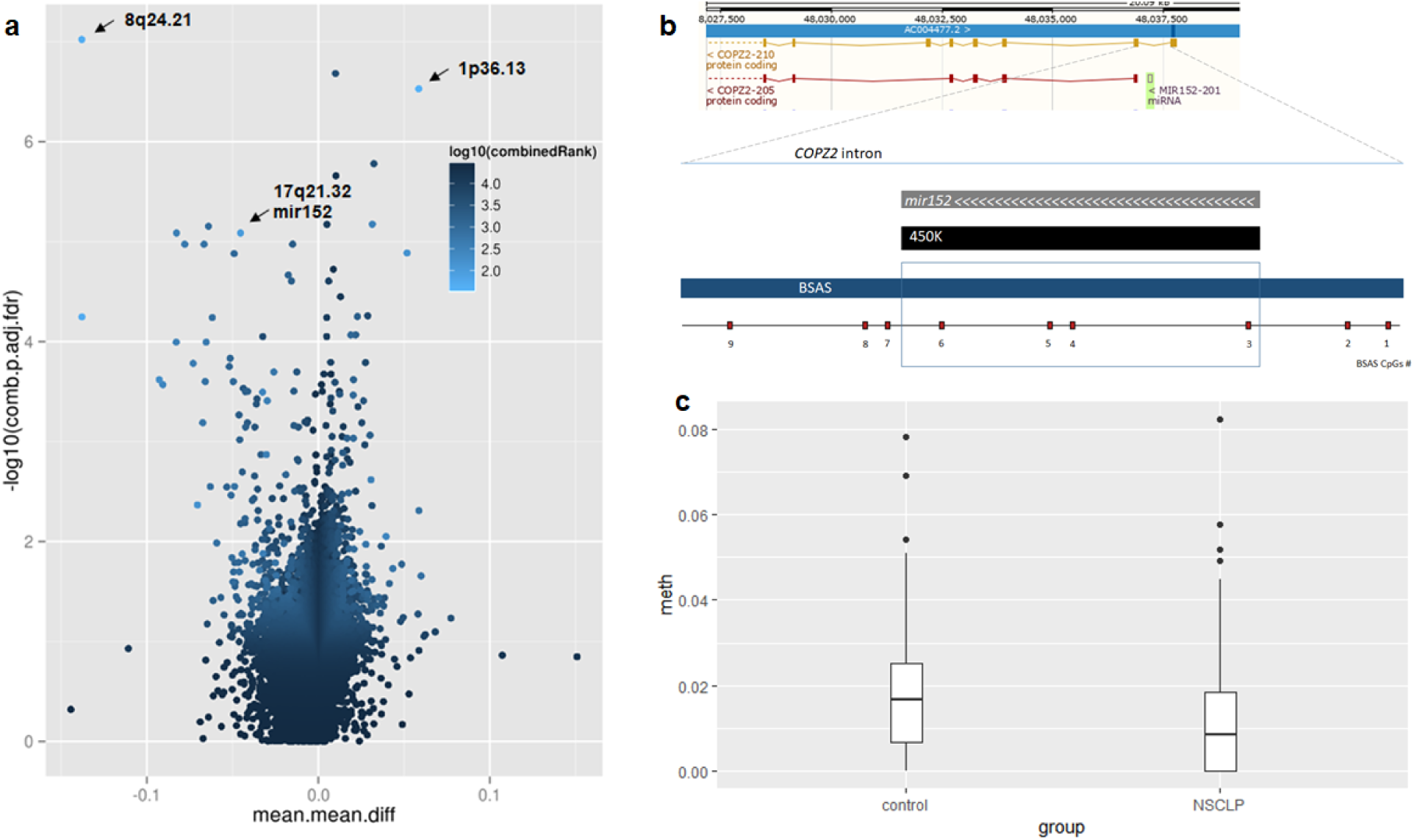
mir152 is differentially methylated in NSCLP cohorts. **a**. Volcano plot of differentially methylated regions (DMRs) at the 450K cohort. Light blue spots are the best ranked DMRs by a p-value, methylation difference and quotient of difference by RnBeads. Arrows indicate DMRs at 8q24.21, 1p36.13 and mir152. **b**. Scheme of mir152 DMR and analysed CpGs at the independente cohort (BSAS – Bisulfite amplicon sequencing cohort). CpGs 3, 4, 5, and 6 are within mir152 gene body and 450K DMR. **c**. mir152 is significantly hypomethylated at the BSAS cohort. Boxplots with central lines as medians. P-value= 0.005 (Mann-Whitney’s test).

*mir152* was the most frequent DMR (n=17 NSCLP samples; ∼26%) with ∼6% of average hypomethylation difference (beta-value reduction) in comparison to controls and was not present in previous published data on common epivariation ^42^.

### mir152 methylation validation in other cohorts

To validate the previous findings, we investigated *mir152*, 8q24.21 and 1p36.13 DMRs in an independent Brazilian cohort of 57 NSCLP samples and 130 control samples, using a different method for DNA methylation quantification (BSAS). 8q24.21 and 1p36.13 DMRs were included in the validation step as both regions have been associated with NSCLP ^7,55,56^. We observed no correlation of potential confounding factors (bisulfite conversion batch, PCR batch, age, sex or origin; Supplementary Figure 1a-e) with BSAS methylation data. Besides, principal component analysis (PCA) did not reveal any evidence of sample stratification which could bias methylation variation in our cohort (Supplementary Figure 1f).

We found no significant methylation differences at either 8q24.21 (average beta-value controls= 0,9792; NSCLP= 0,9725; p=0.41, Mann-Whitney’s test) and 1p36.13 (average beta-value controls= 0,1279; NSCLP=0,1225, p=0.08, Mann-Whitney’s test) DMRs in the replication cohort. However, we found significant hypomethylation at the *mir152* DMR (comprising CpGs 3, 4, 5 and 6) in NSCLP in comparison to controls (p=0,005, Mann Whitney test; Figure 1b-c), corroborating our initial findings. To investigate *mir152* hypomethylation at individual NSCLP samples in this independent cohort, we computed those samples with complete hypomethylation (average beta-values at CpG sites 3, 4, 5 and 6 = 0) (Supplementary Table 1). Considering the *mir152* DMR, we found hypomethylation at 16 NSCLP samples (28%) and 17 control samples (13%), which represents a hypomethylation enrichment of 15% (p=0.02, Fisher’s Exact Test). Also, when we considered each CpGs within *mir152* DMR independently, we found hypomethylation enrichment at CpG 3 (18%), CpG 4 (14%), CpG 5 (16%), CpG 6 (21%) and the adjacent CpG 7 (12%). Correlation analysis of methylation levels from all 9 *mir152* CpGs revealed a trend of hypomethylation shared by CpGs 4, 5, 6 and 7 and mild correlation values (Supplementary Figure 2a-b), which could be indicative of a more cohesive methylation block at those sites. Taken together, our results corroborate *mir152* hypomethylation in both Brazilian NSCLP cohorts.

**Figure 2:**
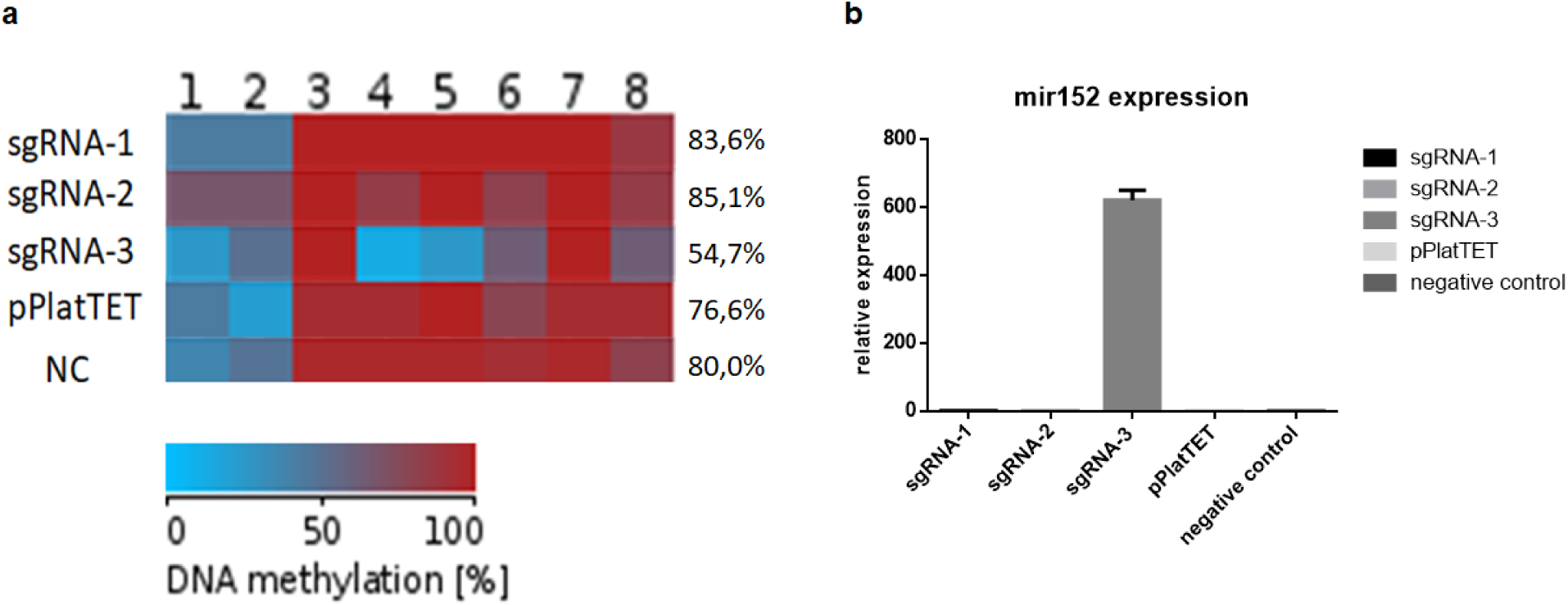
DNA methylation changes at mir152 DMRs results in mir152 expression changes. **a**. A cas9 based approach for target demethylation using the vector pPlatTET and 3 single-guide RNAs sequences (sgRNA 1, 2 and 3) for mir152 DMR in hek293T cells. sgRNA-3 efficiently reduces mir152 methylation, especially at CpGs 4 and 5, in comparison to the empty vector transfection (pPlatTET) and non transfected cells (NC). CpG 9 is absent in hek293T cells due a single nucleotide polymorphism. Total percentage of methylation is represented with values at the right **b**. DNA hypomethylation induces mir152 overexpression in hek293T cells. mir152 expression is absent in all other conditions.

Attempting to evaluate *mir152* methylation contribution to NSCLP in an independent and different population, we looked for other NSCLP methWAS available data. Using summary statistics data from an available NSCLP case-control methWAS performed on 182 hispanic and non-hispanic samples (94 NSCLP and 88 controls ^27^), we did not found significant differences at the *mir152* DMR here studied. On the other hand, we found a CpG site at *mir152* promoter hypermethylated in NSCLP in this cohort (cg06598332, p=0.04), which is located at ∼200bp upstream *mir152* DMR.

### Epivariation is not mediated by genetic variation at mir152 region

Because genetic variation can influence nearby epivariation ^57,58^, we looked for single nucleotide variants (SNV) within the *mir152* DMR. The only polymorphism revealed by Sanger sequencing, rs12940701 (C>T), was present in 30,39% of NSCLP and 41,46% of control samples, with no significant difference between groups (Fisher’s exact test =0,08). Rs12940701 coincides with CpG site 8 at *mir152* DMR, which displays low methylation levels in both NSCLP and control samples (NSCLP average beta-value= 0.0178, controls average beta-value=0.0121). Even though rs12940701 has been suggested as a potential variant diminishing methylation levels at *mir152* region ^59^, we observed no genotype vs. methylation correlation in our replication cohort (p=0.1843, Supplementary Figure 2d). Also, we found no linkage disequilibrium between this SNV and rs1838105, a 1.3Mb apart SNV previously associated with NSCLP at 17q21.32 (Yu et al., 2017; Supplementary Figure 2c). Rare variants (minor allele frequency < 0.5%) at *mir152* gene were not analysed in this cohort (data not shown). Attempting to verify whether rare variants at *mir152* region could segregate in independent NSCLP familial cases, we analysed the exome of 36 affected individuals from 11 families, but neither common nor rare variants were found.

### Methylation variation at mir152 regulates gene expression

We next verified whether methylation variation within *mir152* DMR would be functional and interfere in *mir152* expression. To achieve that, we carried out a CRISPR-Cas9-based approach for targeted demethylation, in which dCas9 were fused to TET1 (pPlatTET-GFP) in order to demethylate specific genomic targets ^48^. Among the three tested sgRNAs (sgRNA-1, sgRNA-2 and sgRNA-3) targeting the *mir152* DMR in hek293T cells, sgRNA-3 efficiently reduced methylation levels at *mir152* DMR at sites 1, 4, 5 and 8 (average beta-value pPlatTET-sgRNA-3 = 0.54, beta-value pPlatTET-NC= 0.76, beta-value NC =0.80; Figure 2a). In non-transfected conditions, or when transfected with the empty vector (pPlatTET-NC) or sgRNAs-1 and 2, hek293T do not normally express *mir152*. On the other hand, consistently with those methylation changes, we observed a high upregulation of *mir152* RNA levels when sgRNA-3 transfections were carried out (Figure 2b). Taken together, those results indicate that epivariation at those sites are functional, resulting in *mir152* expression changes.

### mir152 mimics results in craniofacial malformation in zebrafish

We attempted to investigate whether *mir152* expression could influence craniofacial development. To model that, we injected miRNA inhibitor and mimics in 1-cell stage zebrafish embryos and observed their development at 24hpf and 5dpf. When injected with *mir152* inhibitor, embryos developed normally with no obvious development impairment (Figure 3a). However, when injected with mimics, zebrafish embryos presented several craniofacial defects at 5dpf including malformation of Meckel’s, palatoquadrate, ceratobranchial and the ethmoidal plate, which is the embryo’s analogue palate. Seventy percent (70%) of embryos were affected, which were classified as mildly affected (28,5%), comprising those embryos with ethmoidal plate’s defects in size and shape, and severely affected (41,5%), characterized by a typical cleft at the ethmoidal plate.(Figure 3a). On the other hand, co-injection of *mir152* mimics and inhibitor led to non-affected embryos (n= 65) (Figure 3a).

**Figure 3:**
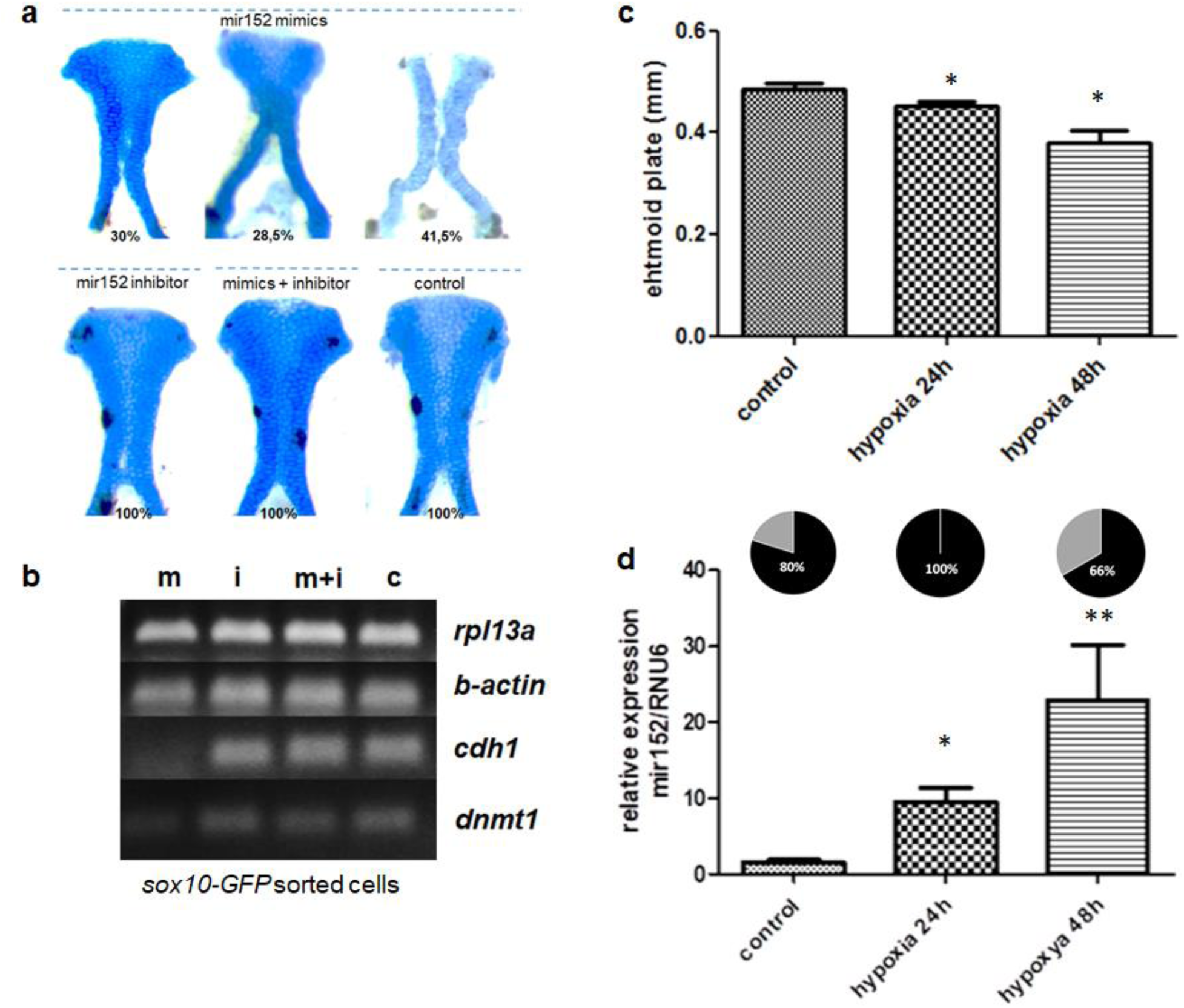
mir152 mimics injected in zebrafish embryos causes ethmoidal plate deffects analogue to clefts. **a**. ethmoidal plates dissected from 5dpf zebrafish larvae injected with mir152 mimics (superior) and mir152 inhibitor, mir152 mimics + inhibiotr and non injected controls (inferior). Mir152 mimics injections resulted in 30% of larvae with non-affected ethmoid plate (left), 28,5% of larvae with mildly-affected ethmoid plate (central) and 41,5% of larvae with severe affected structures, including a cleft ethmoid plate (righ). mir152 mimics injected embryos N= 49. Both mir152 inhibitor injections and mimics + inhibitor combined injections resulted in no altered craniofacial structures with 100% of larvae with normal ethmoid plates. mir152 inhibitor injected embryos N=40; mimics+inhibitor injected embryos N=65. Control embryos N=107. **b**. RT-PCR of cdh1 and dnmt1 in mimics and inhibitor injected embryos. mir152 mimics injection resulted in cdh1 and dnmt1 downregulation. cdh1 expression is not detected in mir152 mimics injected embryos (m) in comparison to mir152 inhibitor injection (i), mimics+inhibitor injection (m+i) and non-injected controls (c). rpl13a and b-actin were used as endogenous controls. **c**. zebrafish embryos in hypoxia (1% oxygen) for both 24h and 48h starting at 2-cell stage presented significant reduction in ethmoid plate’s size (p>0,05, t-test). **d**. mir152 expression significantly increases under hypoxia conditions for 24h or 48h. (*p<0,05; **p<0,005). At the top, pie charts of mir152 methylation levels in zebrafish embryos during normal and hypoxia conditions: controls = 80%, hypoxia 24h = 100% and hypoxia 48h = 66%.

### mir152 targets dnmt1 and cdh1 in zebrafish neural crest cells

Several coding genes are predicted to be targeted by *mir152*, including *DNMT1*, which has been experimentally confirmed ^59–62^. In that case, *mir152* is known to control a *DNMT1/CDH1* loop, in which *mir152* upregulation leads to *DNMT1* downregulation and as a consequence *CDH1* expression is present ^60^. Therefore, we checked for *DNMT1* and *CDH1* expression differences in pPlatTET-GFP transfected hek293T cells or mimics/ inhibitor injected zebrafish embryos. We did not observed *DNMT1/dnmt1* or *CDH1/cdh1* expression changes in any sgRNA transfected hek293T cells or in *mir152* mimics/ inhibitor injected zebrafish whole embryos RNA (Supplementary Figure 3). Because whole embryo expression could mask tissue-specific changes and given the extent of observed craniofacial impairment in injected embryos we hypothesized that neural crest cells (NCCs) would be the most likely cells to be affected by *mir152* dysregulation. To test that, *mir152* mimics/ inhibitor injected embryos were dissociated at 24hpf and sox10-GFP positive were sorted for *dnmt1* and *cdh1* expression analysis. We observed an ablation of *dnmt1* and *cdh1* expression in sox10-GFP positive cells when *mir152* mimics was injected and no effects on *dnmt1* or *cdh1* expression in inhibitor or mimics+inhibitor injections (Figure 3b).

### Hypoxia drives mir152 hypomethylation and expression changes during development and affects craniofacial development

*mir152* expression induction has been reported in cells subjected to hypoxia ^61^ and hypoxia is a known clefting factor in mice and has also been reported to induce ethmoid plate defects in zebrafish ^63–65^. We then hypothesised hypoxia as an environmental factor leading to mir152 hypomethylation, which in turn would cause up-regulation of mir152, resulting in craniofacial malformation. We then first exposed zebrafish embryos to hypoxia (1% O2) for 24h or 48h and obtained 5dpf embryos with reduced ethmoid plate size (Figure 3c). By quantifying *mir152* levels in such conditions, we observed that hypoxia induced a significant up-regulation of *mir152* in comparison to normoxia (up to ∼20 fold, Figure 3d). *mir152* methylation changes in hypoxia were also tested and we observed hypomethylation of *mir152* methylation levels from 80% at normoxia conditions, 100% at hypoxia for 24h and 66% at hypoxia for 48h (Figure 3d). Therefore we consider that hypoxia was capable of reducing *mir152* methylation at least in 14% for a 48h hypoxia treatment.

## Discussion

The study of epivariation on several diseases, especially methylome analysis, have been gaining force in the past five years ^24,25,27,36,37,42,66,67^. In the case of NSCLP, methWASs have demonstrated the association of methylation changes in genes belonging to Epithelial-to-mesenchymal transition (EMT) pathway and also methylation changes associated to cleft subtypes ^24,26^. Whether such epigenetic changes are associated with genetic variation and/or environment is still an open question which we attempted to address in this work. Since environment significantly impacts epigenetic variation ^68^, those findings suggest that, in spite of the high genetic contribution to those phenotypes, environment plays an important role in their aetiology.

By reanalyzing previously published data, we were able to identify *mir152* as a new candidate NSCLP gene in up to 26% of NSCLP samples with DMR hypomethylated. e also confirmed hypomethylation of *mir152* in 15%-21% of NSCLP samples in an independent Brazilian cohort. These findings corroborate our initial findings and suggests a common epivariation at mir152 for NSCLP, which is one of the most frequent alteration so far associated with NSCLP, at least in our population. We also found significant *mir152* promoter methylation differences using methWAS data from a different population ^27^, which suggest that not only epivariation at *mir152* gene body could be associated with NSCLP but also at the promoter region. We believe therefore that epivatiation at *mir152*, both at promoter or gene body, is associated with NSCLP. We could not replicate, however, 8q24.21 or 1p36.13 DMRs in this independent cohort. Methylation differences at 8q24 in NSCLP have been previously reported ^27^, although in a different region than 8q24.21 DMR, more specifically at 8q24.23 *HEATR7A*. Therefore we do not know whether such methylation changes at 8q24 are dependent on each of the studied population or their effects are smaller for detection in our independent cohort.

*mir152* is a member of the mir148/*mir152* family and is located within an intron of *COPZ2* at chromosome 17q21.32, a genomic region previously associated with NSCLP by GWAS, however with *WNT9B* as the principal candidate gene ^7^. Attempting to identify if *mir152* variation could add to the NSCLP GWAS signals at 17q21.32, we also looked at LD data from 1000 genomes and found that both genes were not in LD. Still, we here show that regions previously associated with NSCLP by genetic approaches can also be prone to epigenetic changes as previously reported ^27^.

Because genetic variation can modulate DNA methylation within a region ^57,69,70^, we investigated whether common genetic variation could modulate methylation changes at *mir152*. Our results suggested that neither common nor rare variants at *mir152* region contribute to *mir152* epivariation as a methylation QTL (meQTL) in the cohort here studied. In fact, *mir152* processed sequence is highly conserved and identical from fish to mammals ^71^, indicating that either its function has been conserved during evolutionary diversification and/or genetic variation at that region is not tolerated. We cannot rule out that genetic variation in the promoter region of *mir152* or out of the analysis region, which is not covered in our Sanger sequencing and exome analysis, could lead to expression variability. Further, we also cannot exclude that other tissues than the here studied could display a meQTL status for rs12940701.

To determine the functional effects of *mir152* hypomethylation in gene expression, we induced a cas9-mediated demethylation of *mir152* in hek293T cells, and showed that *mir152* hypomethylation leads to *mir152* upregulation in human cells. Importantly, the major methylation changes were made at CpGs 4 and 5, the core of mir152 DMR, which coincides with the CpG sites bearing higher methylation correlation. Therefore the findings on *mir152* hypomethylation at both 450K and the independent cohorts are likely functional.

Once we found *mir152* hypomethylation to promote *mir152* upregulation, we mimetized *mir152* upregulation in zebrafish development by *mir152* mimics injections. *mir152* upregulation led to several craniofacial defects compatible with clefting phenotypes, grouped in two severity degrees:: mid-affected and severely-affected zebrafish larvae at 5dpf. We speculate that the extension of those phenotypes are dose dependent, because micro-injection of zebrafish embryos can vary in precision of oligonucleotides incorporation by the embryo cells. It is important to note that such observed phenotype were specific to *mir152*-mimics injection, compatible with a *mir152* upregulation scenarium, once both inhibitor and mimics + inhibitor injections resulted in no affected embryos.

Functional studies have demonstrated *mir152* as an important modulator of TGFbeta-induced EMT in epithelial cells, in which *mir152* overexpression is known to inhibit TGFbeta and therefore EMT ^72^. It has also been shown that upregulation of *mir152* targets *DNMT1*, which in turn controls *CDH1* expression via DNA methylation and therefore affecting E-cadherin levels and EMT in breast cancer cells ^60^. Interestingly, *CDH1*/E-cadherin loss-of-function mutations have been found in both syndromic and nonsyndromic clefting forms and *CDH1* promoter hypermethylation have been found in association with cleft penetrance in NSCLP families ^13,24,73^.

In this study, We observed a reduction of both *dnmt1* and *cdh1* expression in *mir152* mimics-injected sox10-GFP NCCs suggesting that this control can be specific at certain cell types, more specifically NCCs. Even more interestingly and opposite to what has been described by other works, which reported mir152 upregulation accompanied by *dnmt1* downregulation and *cdh1* upregulation, we found total reduction of *cdh1* expression when *mir152* mimics was injected. We do not know if this effect is related to *dnmt1* or to other targets of *mir152* in this cell population; nevertheless loss of *CDH1* expression is compatible with *CDH1* related molecular pathology in NSCLP families ^13,73,74^ and *cdh1* downregulation in NCCs has been reported to inhibit NCC migration in Xenopus ^75^. Therefore, given the effects of *mir152* upregulation on gene expression of *sox10*-GFP positive cells our results suggest that aberrant expression of *mir152* during development affects proper craniofacial formation by disrupting neural crest specification/EMT and its derivatives

While the vast majority of studies on NSCLP etiology states the multifactorial scenarium for NSCLP, in which both genome and environment play a role, knowledge on how they interact and which effects NSCLP associated environmental factors have on genome behavior and, more specifically, on the epigenome is scarce. Here we hypothesised that such hypomethylation and consequently upregulation of *mir152* was potentially caused by embryonic hypoxia. Hypoxia is a normal condition during several steps of mammalian development required for proper cell differentiation and progression ^76^, however abnormal oxygen levels below the foetal hypoxia limits can lead to malformations and disease ^77–79^. Regarding oral clefts and craniofacial development, hypoxia has been for a long time demonstrated as a strong risk environmental factor in mice, rat and chicken models ^64,65,80–82^ and also hypoxia-related environmental factor are epidemiologically associated to NSCLP ^16,18,20,63,83,84^. More recently, a hypoxia induced clefting model in zebrafish has been demonstrated ^63^ reinforcing the effect of hypoxia on craniofacial development and supporting our model. In agreement with this study, our hypoxia exposure in zebrafish embryos also resulted in aberrant ethmoid plate sizes and in increased *mir152* expression accompanied by *mir152* hypomethylation at 48h of hypoxia. Our work therefore links an epigenetic alteration in NSCLP to a potential environmental factor, contributing to the multifactorial model proposed to this malformation.

In summary, we demonstrated how individual methylome analysis in NSCLP can indicate individual specific methylation changes potentially relevant to phenotype. In that case, we found *mir152* hypomethylated in 26% of our cohort and replicated this finding in 21% of the cases on an independent NSCLP cohort. Methylation changes at *mir152* result in expression changes and *mir152* upregulation during development leads to impairment of craniofacial development and maternal/foetal hypoxia might be the environmental link leading to *mir152* epivariation. We suggest therefore *mir152* as a novel candidate locus for NSCLP, expanding the current knowledge on NSCLP aetiology and molecular mechanisms.

## Methods

### Ethics

This study was approved by the Ethics Committee of the Instituto de Biociências (Universidade de São Paulo, Brazil). Biological samples were collected after signed informed consent by the parents or legal guardians. All experiments were performed in accordance with relevant guidelines and regulations.

### Affected individuals and controls samples

For methylome analysis, we used previously published and public data ^24^, which briefly consisted of Illumina Infinium HumanMethylation 450K data of blood-derived DNA from 66 cases from non-familial NSCLP individuals and 59 age and sex-matched controls from healthy individuals (hereafter named as “450k cohort”). Our replication cohort consisted of 57 non-familial NSCLP and 130 controls samples which were ascertained either at the Hospital das Clínicas of Universidade de São Paulo (São Paulo, Brazil), Centro de Pesquisas Sobre o Genoma Humano e Células-Tronco of Universidade de São Paulo (São Paulo, Brazil) or during missions of Operation Smile Brazil (Supplementary Table 1). Samples from replication cohort were saliva-derived DNA collected with Oragene (DNA Genotek) and genomic DNA extracted as recommended by the fabricant.

### 450Kmethylome analysis

To identify differentially methylated regions (DMRs) at the gene level in NSCLP samples, we first compared all 66 NSCLP samples versus all 59 controls (450K cohort) using the RnBeads pipeline, which comprises filtering, normalisation and differential methylation steps ^43^. We filtered out probes affected by SNPs, on sex chromosomes, probes with a p-value detection >0.05 (Greedycut) and probes with non-CpG methylation pattern. Data was normalised using the SWAN method. Principal component analysis (PCA) were also performed using R packages in order to identify obvious confounding effects in the 450K cohort. Differential methylation analysis was performed using the RefFreeEWAS method, which corrects p-values for blood cellular contributions, accounting for gene regions. We also used sex, age and probe markers of batch effects as covariates for differential methylation analysis p-value correction as previously described ^24^. We used as selection criteria the 5 top ranked DMRs listed by RnBeads, which ranks DMRs combining adjusted p-values, methylation difference and quotient of difference. As a second step to identify individual contributions to the selected DMRs, we conducted analysis individually comparing each NSCLP sample versus all 59 controls using the same parameters above described. At this phase, =we selected as DMRs those regions with p-value < 0.05 after FDR and covariate adjustment and with at least 5% beta-value difference. We also compared those DMRs with previously published data of frequent and common DMRs ^42^. DMRs were listed by NSCLP sample and we checked for DMRs co-occurring in different NSCLP samples.

### Bisulfite amplicon sequencing of mir152 in the replication cohort

To quantify methylation levels at *mir152*, 8q24.21 and 1p26.13 DMRs in the replication cohort, we used the Bisulfite Amplicon Sequencing (BSAS) method as previously described ^24^. In summary, BSAS relies on bisulfite PCR, library preparation and DNA sequencing with a NGS sequencer ^44,45^. We designed bisulfite-specific PCR primers for those DMRs using the online tool MethPrimer (http://www.urogene.org/methprimer/) with reported recommendations to avoid biased bilsufite PCR amplification ^46^. The predicted amplicons in GRCh37/hg19 build for those DMRs are: mir152 at chr17:46114502-46114660 (Forward sequence: 5’-CS1-GGYGTTGTGTTYGTTGGGTG-3’, Reverse sequence: 5’-CS2-AATCCAACCRGACCAAAAATCAACTA-3’); 8q24.21 at chr8:130876990-130877116 (Forward sequence: 5’-CS1-TATGGAATTGATTAATGAGGAAAAT-3’, Reverse sequence: 5’-AAAACCTTRGATACATTACTAAAAA-3’); and 1p36.13 at chr1:17231171-17231307 (Forward sequence: 5’-GGTGYGTYGAGATTTTGTAT-3’, Reverse sequence: 5’-TTCCAATCTACTATTAAAAACCAT-3’). Samples from the replication cohort DNAs were submitted for bisulfite conversion using 1ug of DNA in the e EZ-96 Methylation Kit (Zymo Research). Converted DNA was used as a template for bisulfite-specific PCR with the HotStartTaq Plus (QIAGen) standard protocol and amplicons were checked by agarose gel electrophoresis and by Bioanalyzer HiSensitivity DNA prior to library preparation. During the library preparation indexes were added in one PCR step for sample (Access Array Barcode Library, Fluidigm). Libraries were purified by Ampure XP Beads in a magnetic column and checked again in the Bioanalyzer HiSensitivity DNA for peak shift visualization. Finally libraries were submitted for sequencing with the MiSeq Reagent V3 Kit 150 bp single-ended run on a MiSeq Sequencer (Illumina). We performed de-multiplexing of sequences using the FASTX Barcode Splitter program in the FastX Toolkit R package (http://hannonlab.cshl.edu/fastx_toolkit/). Following this, we filtered out reads of low quality, selecting only reads with at least 50% of bases with Q > 30 using the FASTQ Quality Filter program, also part of the FastX Toolkit R package. Next FASTQ files were converted to FASTA files using the FASTQ-to-FASTA program in the same package. For the quantification of methylation levels at the mir-152 region we used the BiQAnalyzer HT software ^47^, in which we applied quality filters as follows: minimal reference sequence identity to 90%, minimal bisulfite conversion rate of 90%, maximum of 10% gaps allowed in CpG sites and minimal of 10 reads of coverage. Following these parameters, we obtained average *mir152* region methylation level per sample and also site methylation level within *mir152* region. To investigate hypomethylation, we calculated the controls’ 10th percentile and computed NSCLP samples below this limiar. Frequencies were tested by expected in controls and observed in NSCLP using Chi-square test. Graphs were generated using R package ggplot2.

### Independent population NSCLP methylome data

We used summary statistics data public available from an independent NSCLP case-control methylome study performed on 182 hispanic and non-hispanic individuals ^27^. We looked for significant (p>0.05) probes overlapping *mir152* region (cg02742085, cg05096161, cg05850656, cg06598332, cg09111258, cg10382221, cg10472567, cg21384971, cg24389730).

### Sequencing genetic variation and exome analysis at mir152 region

For sanger sequencing we PCR amplified *mir152* region in replication cohort samples using primers forward 5’-TTCTGGGTCCGTTTGGAGTG-3’ and reverse 5’-TCAAGGTCCACAGCTGGTTC-3’ and Platinum Taq Polymerase Supermix. Amplicons were treated with ExoProStar (GE Healthcare Life Sciences) and then submitted to Sanger sequencing using the BigDye Terminator v3.1 Sequencing standard kit (Applied Biosystems). Next, sequencing products were purified using Sephadex G-50 (GE Healthcare Life Sciences) with MultiScreen Column Plates (Merck-Millipore) and finally submitted to capillary electrophoresis at the ABI 3730 DNA Analyser (Applied Biosystems). All reactions were performed using fabricant’s recommended protocols. Variants in *mir152* were also analyzed in exome sequencing data of 36 NSCLP individuals belonging to 11 families, segregating the disorder under an autosomal dominant model with incomplete penetrance (Supplementary Figure X). Library preparation and exome capture were performed using: Illumina’s TruSeq DNA Sample Prep and Exome Enrichment kits (for families F617, F886, F2570, F3196 and F7614); Illumina’s Nextera Rapid Capture Exome (for families F1843, F2848, F8418), and Agilent’s Sure Select QXT Target Enrichment (F10950, F10955 and F11730). Library quantification was performed with NEBnext Library Quant Kit (New England Biolabs), prior to paired-end sequencing on HiScanSQ (Illumina; families F617, F886, F2570, F3196, F7614) or HiSeq 2500 (Illumina; families F1843, F2848, F8418, F10950, F10955 and F11730) USA). Exome mean coverage per individual was 131x (49 SD). Sequence alignment to the hg19 reference genome, exome indexing, variant calling and variant annotation were performed, respectively, with Burrows-Wheller Aligner (BWA; http://bio-bwa.sourceforge.net), Picard (http://broadinstitute.github.io/picard/), Genome Analysis Toolkit package (GATK, http://broadinstitute.org/gatk/) and ANNOVAR (http://www.openbioinformatics.org/annovar/).

### Site specific demethylation

To functionally investigate the role of methylation variation at the *mir152* DMR, we used a CRISPR-Cas9 based approach in which a plasmid expressing a modified and catalytically inactive Cas9 (dCas9) were fused to the catalytic domain of TET1 with a co-expression system for sgRNA, allowing target specific demethylation ^48^. We obtained plasmid pPlatTET-gRNA2 (#82559) from Addgene. *mir152* specific sgRNAs were designed with CRISPRdirect (https://crispr.dbcls.jp/), named as sgRNA-1 (5’-TCTGTGATACACTCCGACTC-3’), sgRNA-2 (5’-GCTCGGCCCGCTGTCCCCCC-3’) and sgRNA-3 (5’-TGACAGAACTTGGGCCCGGA-3’). sgRNAs were cloned to plasmids as previously published ^48^. All the three plasmid-sgRNA combinations plus empty plasmids were transfected to hek293T cells with SuperFect (QIAgen) following the fabricant’s protocol. After 48h post transfection, cells were checked by fluorescent microscopy to visualize GFP expression and GFP-positive cells were sorted with the BD FACS Aria II and BD FACS Diva software and then pelleted to simultaneous DNA, RNA and protein extraction using TriPrep kit (MN).

### cDNA synthesis and Real time quantitative PCRs

RNA samples were submitted to cDNA synthesis for miRNA using the NCode miRNA First-Strand cDNA Synthesis kit (LifeTechnologies, USA) and recommended protocols. RTqPCR were performed using Fast SYBRGreen MasterMix (Thermofisher) and *mir152* specific primers with NCode miRNA First-Strand cDNA Synthesis qPCR Universal Primer in a fast mode SybrGreen reaction at the QuantStudio 5 (Thermofisher). We used *RNU6B* and *RNU44* as endogenous controls. Relative expression values were calculated using the Delta Delta Ct method as previously reported ^49^. For mRNA cDNA synthesis, we used the same total RNA (1ug) as inputs for the SuperScript IV First-Strand Synthesis System (ThermoFisher) and specific primers for human *CDH1* (Forward Sequence: 5’-CCATTCAGTACAACGCCCAACCC-3’, Reverse Sequence: 5’-CACAGTCACACACGCTGACCTC-3’), *DNMT1* (Forward Sequence: 5’-TATCCGAGGAGGGCTACCTG-3’, Reverse Sequence: 5’-CTGCCATTCCCACTCTACGG-3’) and *TBP* (Forward Sequence: 5’-GTGACCCAGCATCACTGTTTC-3’, Reverse Sequence: 5’-GCAAACCAGAAACCCTTGCG-3’) and *HPRT1* (Forward Sequence: 5’-CCTGGCGTCGTGATTAGTGAT-3’, Reverse Sequence: 5’-AGACGTTCAGTCCTGTCCATAA-3’) as endogenous controls, as well as zebrafish specific primers for *cdh1* (Forward Sequence: 5’-TGTGACTGCAAAGGAGAGGC-3’, Reverse Sequence: 5’-GAGCAGAAGAAGAGCAAGCAATAG-3’), *dnmt1* (Forward Sequence: 5’-TGTTACTTTGGGCAAGAGGAGAG-3’, Reverse Sequence: 5’-AGTGGTGGTGGCTTTAGTCG-3’) and *rpl13a* (Forward Sequence: 5’-TCTGGAGGACTGTAAGAGGTATGC-3’, Reverse Sequence: 5’-AGACGCACAATCTTGAGAGCAG-3’) and *beta-actin* (Forward Sequence: 5’-CGAGCTGTCTTCCCATCCA-3’, Reverse Sequence: 5’-TCACCAACGTAGCTGTCTTTCTG-3’) as endogenous controls, in a SybrGreen reaction at the QuantStudio 5 (Thermofisher) or conventional PCR. For zebrafish *mir152* quantification we used Taqman microRNA assay and probes for dre-*mir152* and rnu6, following the manufacturer’s recommendations.

### Bisulfite sequencing on hek293T transfected cells

For *mir152* methylation analysis after pPlatTET1-GFP plasmid transfections in hek293T cells, we applied traditional bisulfite sequencing method, consisted of bisulfite conversion of 1ug of genomic DNA and PCR amplification of *mir152* region using the method above described. PCR products cloning into pGEM-T-easy vector system (Promega). We Sanger sequenced 10 colonies per sample using M13 primers using the above described method and results were analysed with BISMA online tool (Bisulfite Sequencing DNA Methylation Analysis - http://services.ibc.uni-stuttgart.de/BDPC/BISMA/) ^50^ with default parameters.

### Injection of mirna mimics and inhibitor in zebrafish embryos and hypoxia tests

We performed crossings of both AB and sox10-GFP zebrafish lineages and embryos were collected in E3 medium. Specific Mirna mimics and inhibitor for *mir152* were purchased from mirVana Thermofisher Scientific. Embryos at the 1-cell stage were injected with 2nl of 25uM dre-*mir152* mimics, 25uM dre-*mir152* inhibitor, 25uM dre-mimics + inhibitor or TE. Injected embryos were then raised for up to 5 days in E3 medium at 29oC and 12h/12h light/dark cycle. 24hpf injected embryos were collected for RNA extraction and subsequent cDNA synthesis and RTqPCR following the protocols above mentioned, for confocal microscopy imaging or for cell dissociation followed by GFP-positive cell sorting. Pools of 20 embryos at 24hpf were used for cell dissociation following published methods ^51^ and GFP-positive cells were sorted using BD FACS Aria II and BD FACS Diva software. Larvae at 5dpf were collected and fixed in 4% PFA followed by alcian blue staining for craniofacial cartilages phenotyping using previously published protocols ^52^. To study hypoxia effects on zebrafish embryos, we exposed 1-cell stage zebrafish embryos for 24h or 48h in a 1% O2 incubator (Hera Cell - ThermoFisher).

## Acknowledgements

We are thankful to Dr. Passos-Bueno members for help in discussions and lab organization. We thanks to Patrícia Semedo for helping with cell sorting. This work was supported by FAPESP/CEPID 2013/08028-1, FAPESP 2017/11430-7 (LA), 2016/23648-4 (LAB) and CNPq 305405/2011-5 (MRPB) research fellowships.

## Author disclosure statement

The authors declare that they have no conflict of interest.

